# BIOMODULATORY EFFECT OF LOW INTENSITY LASER (830 nm.) IN NEURAL CELLS 9L/lacZ

**DOI:** 10.1101/2021.04.08.438993

**Authors:** Antonieta Marques Caldeira Zabeu, Isabel Chaves Silva Carvalho, Cristina Pacheco Soares, Newton Soares da Silva

## Abstract

**Background:** Currently, research is advancing low intensity laser (LIL) in central nervous system cells to available the benefits of this therapy in neurological disorders, and research seeks to establish the best LIL protocol in biological processes of neuronal tissue with the different energy wavelengths(λ), and exposure time (s) and frequency(Hz). The aim of this study is to check the biomodulatory effects of the LIL in neural cell culture.

**Methods:** Diode laser λ = 830 nm, power of 40 mW, continuous mode, applied in the cells lineage 9L/lacZ, with energy densities of 0.5, 1.5 and 3.0 J/cm2. Analysed 24 hours after irradiation, the results of the cell viability show the difference between the control and treated groups.

**Results:** In the occurrence of apoptosis, no significant manifestation was observed between the control group compared with the irradiated one (P = 0.9956); there was a significant difference between apoptosis and death by necrosis has been between the control and treated groups (P <0.001). In the comet assay there was not difference.

**Conclusions:** With the aim of evaluating whether LIL promotes early activation of programmed cell death, of 9L/lacZ cells, in the proposed parameters of LIL, we observed that promoted an increase in the number of neural cells, highlighting the action of biomodulation; LIL did not promote the activation of apoptosis and did not any indication of DNA deterioration in the comet assay. The results of this study are indicative that the near infrared laser has a positive interaction with neuronal cells.

## 1. Introduction

Neurological diseases, with different brain damages, degenerative or traumatic injuries, kill 6.8 million people per year, accounting for 12% of global deaths, according to the World Health Organization, 2017 (WHO). The total number of people with dementia is estimated to reach 82 million by 2030 and 152 million by 2050. According to Alzheimer’s Disease International, 2018 part of this increase in the population with degenerative neurological conditions in developing countries is the result of the improvement health conditions and quality of life in these countries.

Currently, research is advancing low intensity laser (LIL) in central nervous system cells in order to assess the benefits of this therapy in neurological disorders such as Alzheimer’s, stroke, ischemia, epilepsy, among others (Oron *et al.* 2007; Gao *et al.* 2010; Hashmi *et al.* 2010; Moreira *et al.* 2011; Liebert *et al.* 2014). However, the effective action of the laser on the nerve cell is not yet established. LIL has been described as an alternative treatment of various inflammatory (Anders *et al.* 2010), degenerative (Hashmi *et al.* 2010; Shen *et al.* 2013), tissue repair (Moreira *et al.* 2011), cell proliferation (Oron *et al.* 2007) processes, among others, mainly because it is not an invasive technique. It is believed that the interaction of light with cellular mitochondria increases the production of adenosine triphosphate (ATP) and therefore promotes cellular processes (Karu 1999; Karu & Pyatibrat 2011; Barrett *et al.* 2013). The limits are not defined; minimum and maximum; energy to be deposited in the cell type that does not adversely affect, for example, causing the promotion of early cell death.

The intense scientific research seeks to establish the best LIL irradiation protocol in biological processes of neuronal tissue, in the face of different energy wavelengths (J/s), (λ), and exposure time (s), frequency modulation (Hz), among other parameters.

Many studies are being developed in in vivo experimental models, however there are few results found in neural and neuronal cell extracts. Therefore, it is hypothesized that low-intensity laser irradiation promotes cell biomodulation with cell cycle promotion and other mechanisms involved in cell viability.

The present work objective is to verify if there is early activation of programmed cell death, apoptosis and if the genetic material suffers any deleterious action by irradiation in 9L/lacZ cell at energy densities of 0.5 J/cm^2^; 1.5 J/cm^2^ and 3.0 J/cm^2^ with infrared diode laser.

## 2. Materials and Methods

### 2.1. Low Intensity Laser

The equipment laser used in this study was an ArGaAl diode (DMC®) detailed in Table 1. Equipment’s power was previously verified against a power meter, brand Low pot - Medium Power Thermal Sensors, model 30 (150) A-BB-18 (Ophir Phothonics®), with aperture of 35mm. The irradiation of the cells occurred 24 hours after plating in darker conditions to avoid interference from other light sources. The laser probe was fixed and kept in contact with the lower face of the plate (Figure 1). The irradiation parameters are described in Table 1.

**Figure 1.**
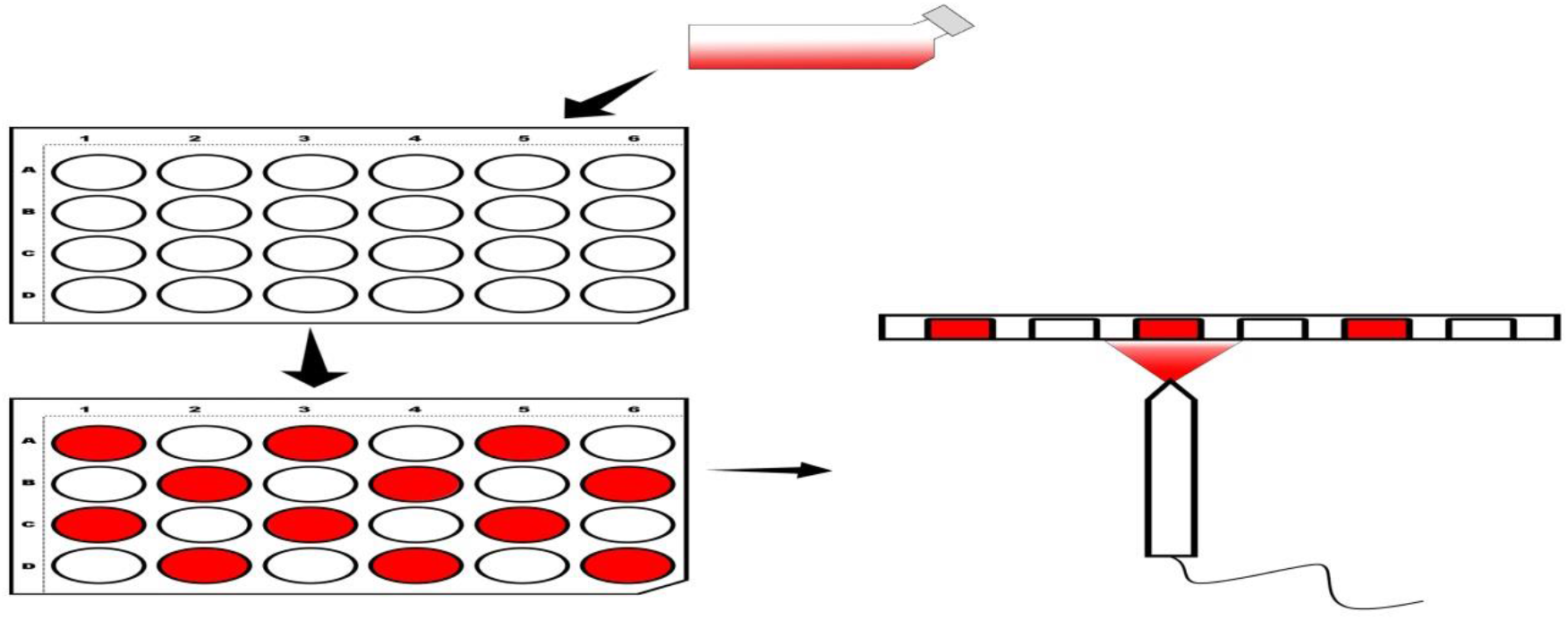
The laser probe was maintained fixed and in contact with the plaque, at the lower surface of the cell container.

**Table 1.** Specification of the equipment used

| Parameters | Laser |
| --- | --- |
| $\lambda$ (nm) | 830 |
| ∅ (mm) | 0.8 |
| P (W) | 0.04 |
| A (cm<sup>2</sup>) | 1.93 |
| mode | continuous |

### 2.2. Cell Culture

The cells lineage employed in this work is the 9L/lacZ (ATCC® CRL2200TM). The cells were cultured in DMEM high glucose (Sigma-Aldrich Co. LLC) supplemented with 10% bovine fetal serum and 1% antibiotics. After a seven day culture in an incubator (CO_2_ Incubator - COM-170AICUV - Panasonic Healthcare) at 37 °C, 5% CO_2_ enviroment, cells were then trypsinized (0.25% trypsin), proceeded to centrifugation (Centrífuga Excelsa Baby I – Fanem - Brasil) at 8000 rpm during 5 min. The cell pellet was resuspended in 2 ml of cell culture medium. An aliquot of the cell suspension was stained with Trypan Blue (10 μL), then this portion was transferred to an automatic counter Countess FL II Automated Cell Counter (Thermo Fischer Scientific Inc.). The automatic counter provided the amount of viable cells, and this information was initially adopted as the amount of cells to be placed in each well for irradiation. Cells were cultured in 24-well polystyrene plates with cell density of 1×10^5^ cells/well. The culture was maintained in an incubator at 37 °C and an atmosphere of 5% CO_2_ for cell adhesion. Four experimental groups were defined: G1 control group, G2, G3, G4 irradiated groups as described in Table 2. For irradiation, the culture medium was removed and replaced with phosphate buffered saline (PBS), avoiding an interference of phenol red staining of the culture medium. At the end of the irradiation, the plate was protected from the action of any other light source. The PBS was replaced by the culture medium and the samples were kept in an incubator at 37 °C in an atmosphere of 5% CO_2_ for 24 hours.

**Table 2.** Energy deposited in each experimental group

| Groups | Energy (J) | Energy Density (J/cm<sup>2</sup>) | Time (s) |
| --- | --- | --- | --- |
| G1 (control) | 0.0 | 0,0 | 0,0 |
| G2 | 0.9 | 0.5 | 24,12 |
| G3 | 2.7 | 1.5 | 73,5 |
| G4 | 5.4 | 3.0 | 144,75 |

Table 2. Energy deposited in each experimental group
| Groups | Energy (J) | Energy Density (J/cm<sup>2</sup>) | Time (s) |
| --- | --- | --- | --- |
| G1 (control) | 0.0 | 0.0 | 0.0 |
| G2 | 0.9 | 0.5 | 24.12 |
| G3 | 2.7 | 1.5 | 73.5 |
| G4 | 5.4 | 3.0 | 144.75 |

### 2.3. Cell Death - Apoptosis - Annexin V

To assess whether the laser induced apoptosis in glial cells, Vybrant® Apoptosis Assay Kit Test # 6 (InvitrogenTM) was used. 24 hours after irradiation, the culture medium was removed and cells were washed with phosphate buffer saline (PBS). The cells were removed with tripsin and centrifuged 8000 rpm for 7 minutes. After centrifugation, the cells were resuspended in 100μl of the staining solution (25μl of Annexin; 5 μL of propidium iodide and 470 μL buffer). For quantitative analysis of apoptotic cells the Tali®-based Image Cytometer (Life Technologies Corporation), based on fluorescence assay, was used. The assay result was shown quantitatively in percent of the apoptosis occurrence in a sample of 25 μl of 9L/lacZ cells, 24 hours after irradiating with low level laser.

### 2.4. Cellular Viability– Violet Crystal

This assay evaluates the cell density, being reflected by the colorimetric determination of cells stained with crystal violet (CV). After 24 hours of irradiation, the culture medium was discarded from each well, 300 μl of staining solution (5% VC, 1.7% NaCl, 3.3% paraformaldehyde, ethanol 33,3% and 56.7% water) were added to each well and incubated for 5 min at room temperature. Cells were then washed in sequence and 300 μl of DMSO (Dimethylsulfoxide) was added to each well and incubated for a period of 1 hour at room temperature. Absorbance of each sample was determined on an ELISA reader Spectracount (Packard Instrument Company Inc., Meriden, CT, USA) with a 570 nm filter. The experiment was performed in triplicate (at different times).

### 2.5. Comet assay

Twenty-four hours after irradiation, 200 μl of trypsin/0.25% EDTA (Gibco) were added to each well for 3 min to detach the cells. After centrifugation (3000 rpm/5 min), each group of cells was resuspended in 200 μl of 0.5% low-melting point agarose (Gibco) and transferred to slides prepared with 1.5% agarose (Gibco). The slides were immersed in cold (4 °C) lysis solution (NaCl 2.5 M, EDTA 100mM, Tris 10 mM/ 1% Triton X-100, 10% DMSO) and kept at 4 °C for 1 h. The prepared slides were placed in a horizontal electrophoresis cube, which was then completed with freshly made alkaline buffer at 4 °C (300 mM NaOH and 1 mM EDTA, pH 13). Electrophoresis was performed at 300 mA and 25 V for 30 min. The slides were removed, washed three times (5 min each) with 0.4 M TRIS–HCl, pH 7.5. 30 μl of ethidium bromide (20 μg/mL) were added to stain the slides, and the observations were made with a fluorescence microscope Leica DMLB and images will be captures via video camera digital Leica DFC 300FX at 200 X magnification. Thirty randomly selected fields were photographed and analysis was performed on 250 randomly selected cells per group (Lovell *et al.* 2008). The images were analyzed in the OpenComet software program, in accord with the comet assay protocol was performed by Carvalho et al. (2016). All steps of these analyzes were performed without direct light to avoid further damage to the DNA of the cells.

### 2.6. Statistical Analysis

The results were in triplicate, statistically analyzed by GraphPad Prism 6.0 (GraphPad Inc., La Jolla, CA) by ANOVA nonparametric test and Tukey’s multiple comparisons test, posttest at 5% significance level.

## 3. Results

The results of the cell viability test, by staining with violet crystal, showed a significant difference regarding the density of viable cells in the culture tested (P <0.001) between the control and treated groups, indicating that there was cell proliferation. Among the treated groups, comparing the 0.5 J/cm^2^ group to the 3.0 J/cm^2^ group, and the 1.5 J/cm^2^ group to the 3.0 J/cm^2^ group, the difference was highly significant (P <0.001). However, when comparing the 0.5 J/cm^2^ group to 3.0 J/cm^2^ group, the statistical difference was lower (Figure 2).

**Figure 2.**
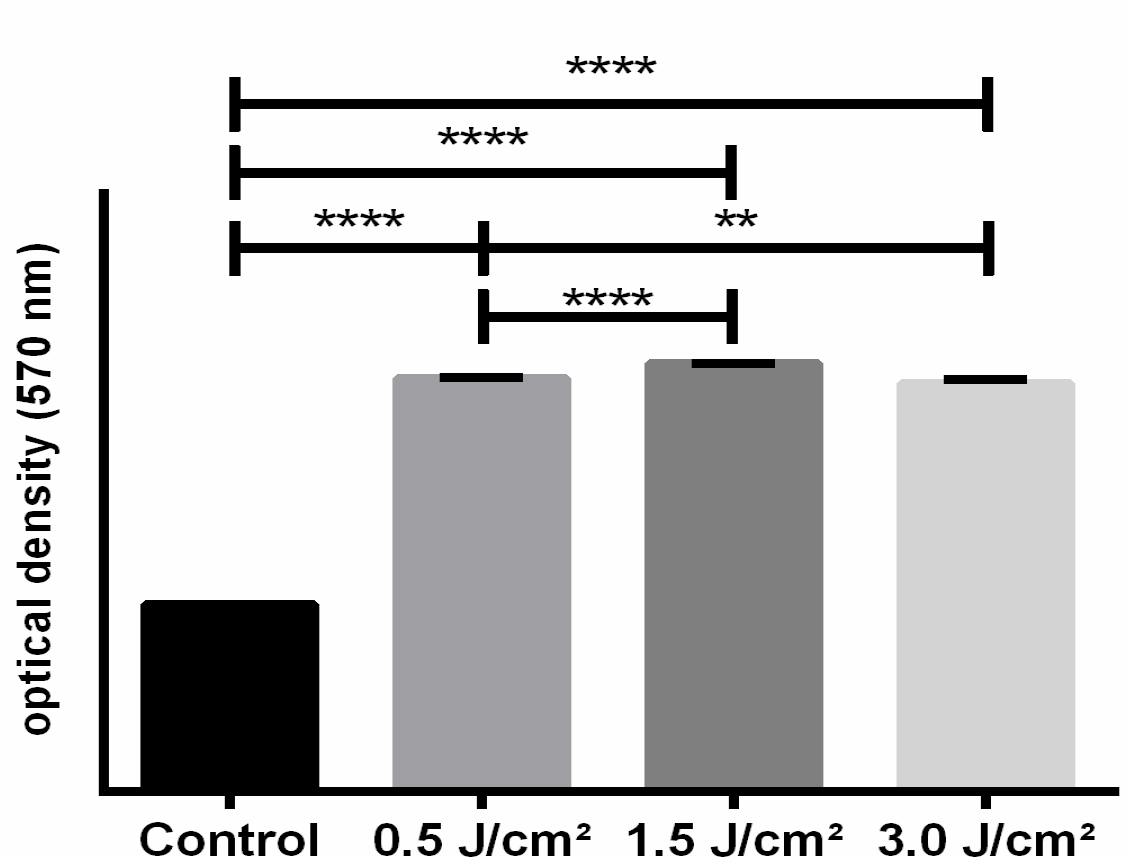
The graph shows the optical density at 570 nm of the viable cells by testing the violet crystal. There is significant difference (P <0.001) between control and treated groups among the groups treated by 0.5 J/cm^2^ with the group of 3.0 J/cm^2^; and comparing groups of 1.5 J/cm^2^ and 3.0 J/cm^2^, there was also a difference in the optical density of irradiated cells.

In the evaluation of live, apoptotic and necrotic cells, using the Annex V assay, the percentage of live cells was considerably high when comparing the groups treated with the control group (P <0.001). In addition, the statistical results show a difference between the groups treated with 1.5 J/cm^2^ and 3.0 J/cm^2^ (P < 0.0093). As for the occurrence of apoptosis, also in percentage, no significant manifestation of programmed cell death was observed when the control group was compared with the irradiated groups (P = 0.8894), nor in the comparison between the treated groups (P > 0.9999). For the evaluation of the percentage of cells killed by necrosis, there was a significant difference in the comparison between the control group and the irradiated groups (P < 0.001) (Figure 3).

**Figure 3.**
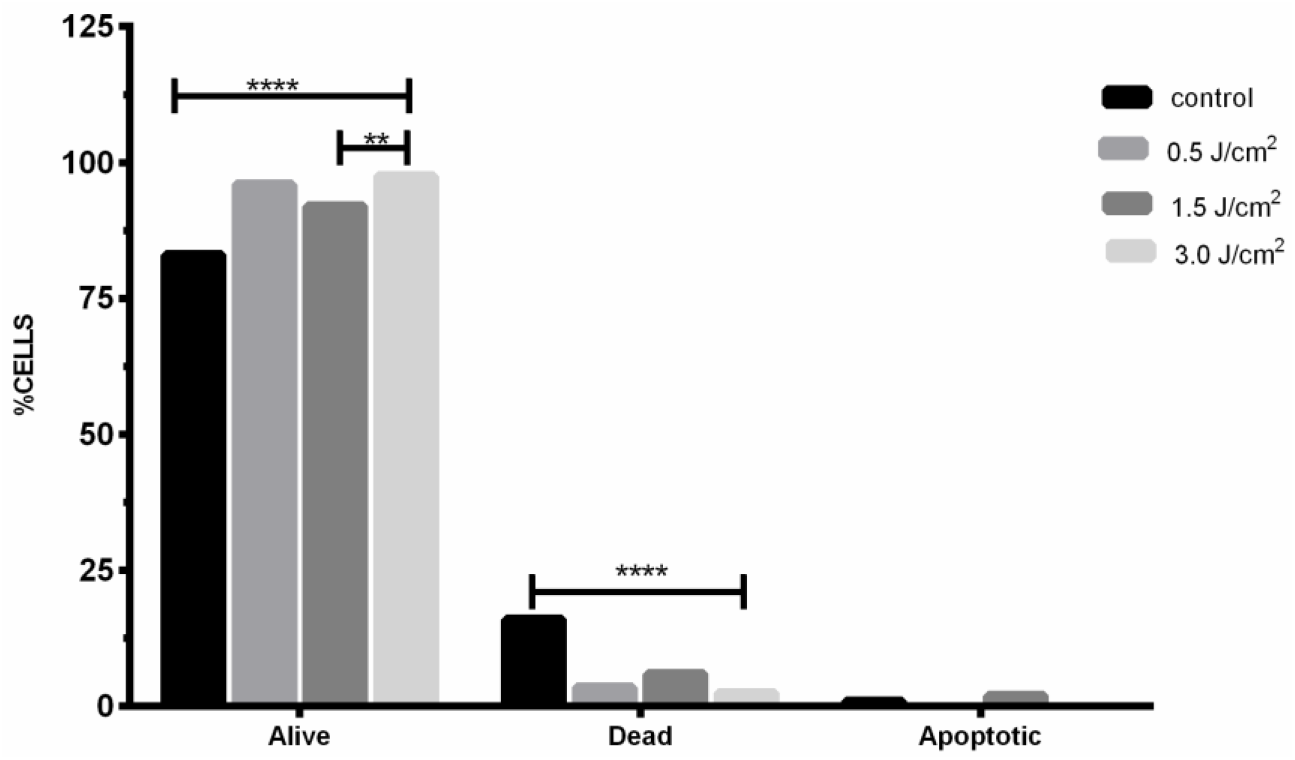
The graph shows the percentage of living cells, and apoptotic cell death, available by Annexin V. The differences were highly significant compared to control treated groups (P <0.001) between the groups treated with 1.5 J/cm^2^ and 3.0 J/cm^2^ (P <0.0093). In the assessment of cell death by necrosis, when comparing between the control group and the treated group, there was no statistical difference observed (P <0.001). The evaluation of the occurrence of apoptosis between control and treated groups (P = 0.8894) and between treatment groups (P > 0.9999).

Average tail length and tail time – taken in arbitrary units – and percentage (%) of DNA in the comet’s tail were used in the results of the comet assay evaluation. (Figure 4) There was no significant difference observed between the treated and control groups, (P < 0.005). There was differenced with little significance in the 0.5 J/cm^2^ group, compared to 1.5 J/cm^2^ (P = 0.0520) in the assessment of% DNA in the comet’s tail. (Figure 3). Thus, the images show the illustration of the cells considered in the Comet assay within the evaluated groups. (Figure 5)

**Figure 4.**
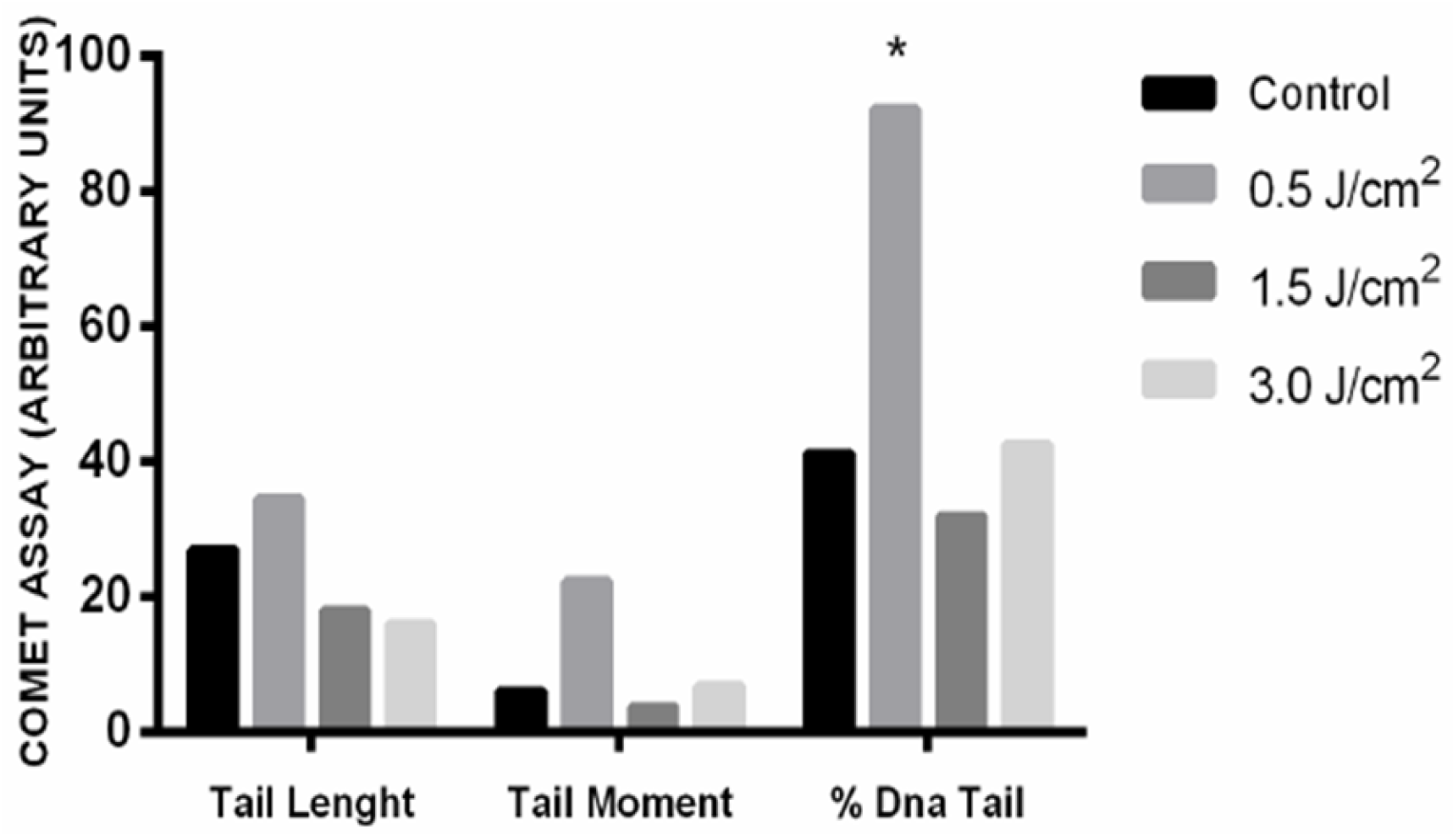
The graph shows length, moment (in arbitrary units) and % DNA in the tail of the comet. No significant difference in the comparison between the treated and control groups has been detected, and there was difference with little significance in the 0.5 J/cm^2^ group, compared to 1.5 J/cm^2^ (P = 0.0520) in the % DNA tail.

**Figure 5.**
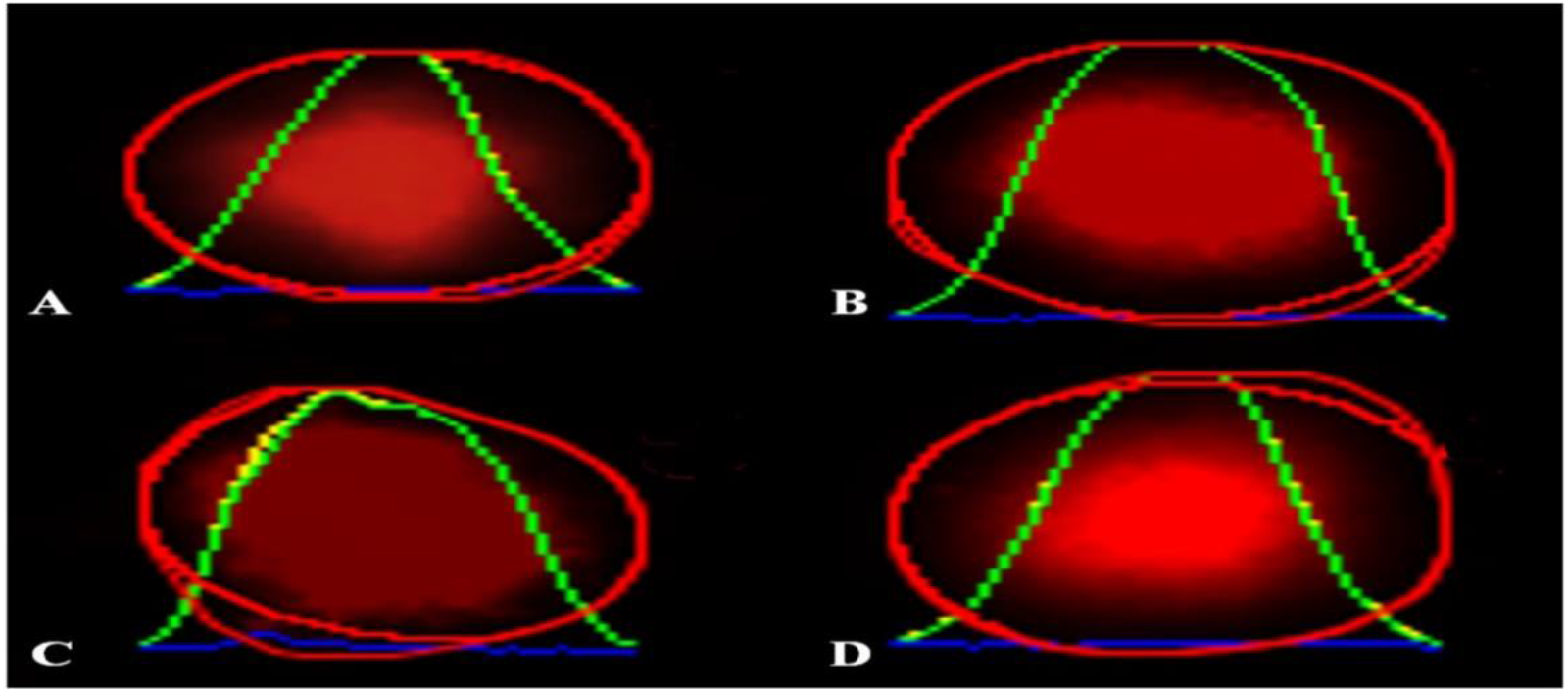
Illustration of cells considered in the Comet assay: (a) 0.5 J/cm^2^, (b) 1.5 J/cm^2^, (c) 3.0 J/cm^2^, (d) no irradiation.

## 4. Discussion

The energy densities used were selected (Mochizuki-Oda *et al.* 2002; Giuliani *et al.* 2009), wavelength (Oron *et al.* 2007; Wu *et al.* 2009) and mode of energy delivery (Wu *et al.* 2009) in different published studies, using infrared lasers on nerve cells (Von Leden *et al.* 2013; Murayama *et al.* 2012). The action of low-power laser in the cell has its mechanism based on the absorption of photon energy by mitochondria cytochrome C oxidase (Gao *et al.* 2009; Barrett *et al.* 2013; Wang *et al.* 2015). This photon receptor plays an important role in neuronal physiology, acting directly on energy metabolism and stimulation of cell signaling pathways; and that this absorption of light by the mitochondrial cytochrome is probably linked to several neuroprotective effects promoted by LIL. In addition, the increase in cellular ATP (adenosine triphosphate) improves the energy support of cells and, consequently, increases cellular processes in general. It has been shown that the increase in adenosine triphosphate is due to the increase in extracellular ATP for dermal cells when irradiated at 657 nm, continuous 35 mW, 0.28 W/cm^2^ at a dose of 17-85 J/cm^2^ over time, from 1 to 5 minutes, hat the aforesaid event occurs that laser promotes positive regulation of ATPase (Rojas *et al.* 2013; Gonzalez-Lima *et al.* 2014; Barboza *et al.* 2014).

According to Hamblin et al. 2016, the effects can be divided into three fronts of action, beginning with the immediate effect of increasing ATP and improving cell oxygenation, followed by activation in the regulatory proteins of anti-apoptotic processes and the production of antioxidants substances such as reactive oxygen species (ROS); finally, there is the formation of neurotrophies and neurogenesis, which both act in tissue repair.

The results of the cell viability test, by staining with violet crystal, showed a significant difference (P <0.001) between the control and the treated groups, regarding cell viability. Among the treated groups, when compared to the 0.5 J/cm^2^ group with the 3.0 2 2 2 J/cm^2^ group; and comparing groups of 1.5 J/cm^2^ and 3.0 J/cm^2^, the difference was highly significant. However, when comparing the 0.5 J/cm^2^ and 3.0 J/cm^2^ groups, the statistical significance was lower (Figure 2). These results clearly show the biomodulator effect of laser therapy in these cells cultures – the maintenance of cell viability is closely linked to the cell’s energy production. The photochemical effects promoted by the interaction of light with mitochondrial cytochrome C and with cellular water triggers a series of biological responses, these being mediated by the production of free radicals that activate cellular mechanisms; as well as an increase in protein synthesis and cytokines that impact cell proliferation (Kim *et al.* 2011; Barolet *et al.* 2016; Tsai *et al.* 2017). The difference between the energy densities of 0.5 J/cm^2^ and 3.0 J/cm^2^, in which the statistical significance was lower in relation to the other groups tested, corroborates the findings of Lubart *et al*. 2006, who reports an additional increase in dose of light energy generates a protective antioxidant effect of the cell. Still, Sommer *et al.* 2012, in their study with neural cells, human neuroblastoma demonstrated that laser prevents oxidative stress and improves mitochondrial efficiency that provides energy for cellular metabolism in general.

Regarding the occurrence of cell death, our study shows a significant difference in the occurrence of death by necrosis when compared to the groups treated with the control (P <0.001) only (Figure 3). This result of death by necrosis may be due to the oxidative stress caused to the cell by LIL, although with statistical significance when we look at the graphic expression of this result, it is clear that the amount of necrotic cells in the sample is small in relation to the resulting proliferation and concentration treatment with LBI. Apoptosis death was not significant in any of the treated groups.

These results suggest that LIL promoted cellular processes, characterizing that there was a biomodulatory effect of the tested cell and increased cellular ATP. A next step in our research project will be the quantification of ATP, ATPase in this experimental model of the 9L/lacZ cell, in order to confirm the hypothesis that LIL increases the energy contribution of this cell with the consequent promotion of its metabolism.

A study comparing two wavelengths, 830 nm and 652 nm, 4.8 W/cm^2^, in continuous mode, directly irradiated in rat brains (Wang *et al.* 2015) showed significant differences in the increase in ATP in the 830 nm laser group. Another report (Sharma *et al.* 2011) using 810 nm, power density of 25 mW/cm^2^ at energies of 0.03; 0.3; 3; 10, 30 J/cm^2^ in continuous mode quantified reactive oxygen species, increased nitric oxide, changes in mitochondrial membrane potential, increased intracellular content of cellular and cellular ATP.

The wavelength used in this study (830 nm) is selected to be within the infrared spectrum in accordance with the literature search. In addition, infrared experiments reported by Von Leden *et al.* 2013 (808 nm, 50 mW, 0.2, 4, 10 and 30 J/cm^2^) and Murayama *et al.* 2012 (λ 808 nm, 30 mW, 18, 36 and 54 J/cm^2^) demonstrated that the interaction of neuronal cells is better at infrared wavelengths. The literature confirms the results found in this study, which used infrared laser, λ 830 nm.

Some authors (Karu 1999; Karu & Pyatibrat 2011; Sharma *et al.* 2011) describe that the increase in ATP and HeLa cell cultures increases cell responses depending on the proliferative phase in which the cell is demanding more or less energy consumption. Carnevalli *et al.* 2003, demonstrated that LIL improves the cell division process, keeping the membrane and the genetic material preserved after irradiation with ovarian epithelium from the λ 830 nm, 10 mW, 2 J/cm^2^ cell culture. In addition, other authors (Von Leden *et al.* 2013; Sharma *et al.* 2011; Evans *et al.* 2009; Yazdani *et al.* 2012) in their experimental studies showed that results such as cell proliferation depends on fluency, when it is very high promoting an opposite effect of inhibiting cell proliferation. The results obtained (Figure 4) showed that the cells even at a dose of 3.0 J/cm^2^, which was the higher dose in this protocol, maintained its entire preserved genetic material by supplementing the information obtained in test Annexin V, described in the preceding paragraphs, where there was no apoptotic death and cell proliferation. These results corroborate the literature and the comments we obtained in the essay of the comet in which we carried out this experiment, as discussed in sequence.

The technique chosen to assess the occurrence of DNA fragmentation in this experiment was electrophoresis in microgel cells, commonly called a comet assay, used to assess DNA damage and cell repair, individually, cell by cell, and consists of lysis of cell membrane, followed by the migration of the DNA released in the agarose matrix. When viewed under fluorescence microscopy, the shape of the cell has the appearance of a comet, where the head is the nuclear region and the tail is the fragments or strands of the damaged DNA. The comet assay is part of the International Organization for Standardization (ISO), number 10993-3-2014, to assess genotoxicity; it is a test that clearly demonstrates the occurrence of damage to cellular DNA, no doubt. It is a test widely used to investigate the genotoxic potential of highly toxic substances, such as heavy metals, pesticides, anti-tumor drugs, among others (Moller 2018).

In addition, this test is a good tool in the investigation of cell repair, in conjunction with studies of apoptosis and cell necrosis –- just as it was done in this study. As already discussed in the previous paragraphs, our tests for evaluating the viability of the cells of this experimentation and programmed cell death showed that biomodulation occurred and already signaled the probability that there would be no damage to the genetic material of these cells. Then, our results, in the evaluation of the comet test, showed that the low intensity laser, within this experimental protocol, with the parameters used for this experimental test, did not cause cell DNA fragmentation after 24 hours of contact. The study performed by Kong *et al*. 2009, in which the different responses of HeLa cells irradiated with different wavelengths, between ultraviolet and near infrared, and which induced lesions directly in the cell nucleus have been analyzed in vitro, has shown that double-stranded DNA disruption can occur at all laser wavelengths, with blue and green lasers causing more aggressive damage to cellular DNA. However, the results of cells irradiated by infrared laser (using sapphire laser at 200 fs, 800 nm, continuous) showed damage to the genetic material. Nevertheless, they observed that DNA repair was promoted quickly, and the injured area was smaller, without propagation and being sensitive to the detection of the comet test. The authors concluded that damage to cellular DNA is directly dictated by the applied energy, wavelength, duration of irradiation and pulse rate as well as the cellular damage repair mechanism, thus corroborating the results that we could observe in our study. This behavior of the 9L/lacZ cell against low-level laser irradiation is possibly due to the activation of the cell’s protection mechanisms, anti-apoptotic proteins and neurotrophins, which are activated by oxidative stress caused by photons, followed by increased mitochondrial membrane potential, increased O2 circulation and cellular metabolism, resulting in greater production of mitochondrial ATP.

Reynolds *et al.* 2013 reviewed the DNA repair dynamics evaluation studies in detail, in which the induction of cell DNA damage was caused by laser irradiation and the authors demonstrated that both the damage to the cell’s genetic material and the repair of that damage are related to the shape of wave used and where the damage occurs in cellular DNA. Cell damage, exogenous or endogenous, when it exceeds the cell’s repair capacity, causes cell death or mutations that can be incorporated into the cell genome.

When the neuronal cell undergoes the action of photons from the laser light, an excitatory photophysical effect of the molecules takes place and, that is resulted, photochemical effects occurs, such as increased ROS, leading to mitochondrial signaling that activates cytoprotective mechanisms. Believed, also, that not us observing lesions in the genetic material of the tested cells, since cell repair was effectively activated to treat any disorder that the cells suffered from the action of the challenged LIL sample.

One of the objectives of this study is to evaluate whether, within the laser parameters applied to the 9L/lacZ culture cells, it would lead to any disturbance occurring in the cellular DNA by the comet test. However, in this study, 24 hours after irradiation, in evaluating cells for the occurrence of DNA fragmentation, the results showed that in the energy density in the employed range (0.5, 1.5 and 3.0 J/cm^2^); it did not cause any damage to the DNA of glial cells, confirming the results reported in the literature (Kong et al. 2009; Zabeu & Pacheco-Soares 2015).

Even so, the analysis of the results suggests the expansion of the studies, through the investigation of greater precision in the in vitro application of the dose of energy supported by the neural cells (DL-50), looking for similarity with the reports of the scientific literature, which apply the LIL in vivo, in order to establish from which power densities, they start to be harmful, as well as an evaluation of the ideal way to supply this energy, in a pulsed or continuous wave.

## 5. Conclusion

Exist a lot of research and clinical use of low intensity laser (LIL) for neurological disease but no protocol specific for each disorder are discuss. In the present study, we hypothesized that the interaction of laser light at energy densities of 0.5, 1.5 and 3.0 J/cm^2^ with 9L/lacZ cells could lead to some cellular disorder, such as decreasing cell viability, promoting programmed cell death (apoptosis) or necrotic death; or any imbalance and changes in cellular DNA.

Regarding the objective of evaluating whether LIL promotes early activation of programmed cell death, apoptosis of 9L/lacZ cells at different energy densities of the infrared diode laser, we observed that the use of low-power laser promoted an increase in the number of cells, suggesting that the biomodulatory action of the laser. In addition, LIL did not promote the activation of programmed cell death - apoptosis. The results of this study are indicative that the near infrared laser has a positive interaction with neuronal cells.

## 6. Ethics approval and consent to participate

This article is not involved in personal information and ethical approval is not required.

## 7. Consent for publication

Not applicable.

## 8. Availability of data and material

No additional supporting data in this article.

## 11. Acknowledgements

We thank Ph.D. Rafael da Cruz Ribeiro Berti – Laboratory of Environmental and Thermal Engineering – USP/Sao Paulo/Brazil for support in laser diode ArGaAl (DMC®) calibration. Ph.D. Clayton Barcelos Zabeu – Mauá Technology Institute – Sao Paulo/Brazil for support in statistical analysis.

## 12. Authors’ contributions

AMCZ conceived and designed the experiments, analyzed and interpreted data, performed experiments and wrote the manuscript; ICSC assisted in the performed experiments; CPS analyzed and interpreted data; NSS revised the manuscript and gave the final approval of the version to be published. The author (s) read and approved the final manuscript.

